# The annotation and analysis of complex 3D plant organs using 3DCoordX

**DOI:** 10.1101/2021.11.20.469364

**Authors:** Athul Vijayan, Soeren Strauss, Rachele Tofanelli, Tejasvinee Atul Mody, Karen Lee, Miltos Tsiantis, Richard S. Smith, Kay Schneitz

**Affiliations:** Plant Developmental Biology, TUM School of Life Sciences, Technical University of Munich, Freising, Germany; Department of Comparative Developmental and Genetics, Max Planck Institute for Plant Breeding Research, Cologne, Germany; The John Innes Centre, Norwich Research Park, Norwich, United Kingdom

## Abstract

A fundamental question in biology concerns how molecular and cellular processes become integrated during morphogenesis. In plants, characterization of 3D digital representations of organs at single-cell resolution represents a promising approach to addressing this problem. A major challenge is to provide organ-centric spatial context to cells of an organ. We developed several general rules for the annotation of cell position and embodied them in 3DCoordX, a user-interactive computer toolbox implemented in the open-source software MorphoGraphX. It enables rapid spatial annotation of cells even in highly curved biological shapes. With the help of 3DCoordX we obtained new insight by analyzing cellular growth patterns in organs of several species. For example, the data indicated the presence of a basal cell proliferation zone in the ovule primordium of *Arabidopsis thaliana*. Proof-of-concept analyses suggested a preferential increase in cell length associated with neck elongation in the archegonium of *Marchantia polymorpha* and variations in cell volume linked to central morphogenetic features of a trap of the carnivorous plant *Utricularia gibba*. Our work demonstrates the broad applicability of the developed strategies as they provide organ-centric spatial context to cellular features in plant organs of diverse shape complexity.

## Introduction

It remains a salient challenge to understand the generation of biological shape. Gaining comprehensive insight into the multi-scale processes underlying morphogenesis critically depends on the quantitative description of molecular, cellular, and tissue-level parameters, such as gene and protein expression, cell geometry, and cell topology [1–6]. Moreover, as prominently proposed by D’Arcy Thompson, the generation of shape can be achieved by growth that is oriented relative to a coordinate system imposed on the organ [7]. Thus, insight into tissue morphogenesis further relies on putting cellular data into context by placing them within an organ-related frame of reference [8–11].

Realistic 3D digital organs with cellular resolution have become indispensable tools for the study of morphogenesis. They can be obtained by deep imaging of fluorescently marked specimens using for example confocal laser scanning microscopy (CLSM) or light sheet fluorescence microscopy (LSFM) followed by 3D cell segmentation of the obtained z-stacks of optical sections with the help of constantly improving software [11–17]. Tissues and organs of model plants are particularly well suited for the generation of such 3D digital representations. Plant cells are immobile simplifying the detection of cellular growth patterns associated with tissue formation. In addition, plant tissues are characterized by a small number of different cell types and often exhibit a well-structured, layered organization. Thus, they usually feature a cellular anatomy of manageable complexity. Accordingly, a growing number of realistic 3D digital tissues with cellular resolution are being generated, mainly in the model plant *Arabidopsis thaliana* [9,10,18– 27].

With the help of 3D digital organs quantitative information about geometric and molecular parameters of up to thousands of cells can readily be obtained using open-source software, such as MorphoGraphX [11, 14] (preprint). However, meaningful exploration of such complex data sets remains challenging. In particular, it is important to provide spatial context by placing the cell’s data within an organ-related frame of reference [8]. Several computational pipelines have been established that provide such a tissue-level frame of reference and allow the semi-automatic annotation of 3D cellular properties in a plant tissue context with cellular resolution: iRoCS [9], 3DCellAtlas [10], and 3DCellAtlas Meristem [22]. These computational pipelines have been applied very successfully for the annotation of cells and tissues in the main root and the hypocotyl, radially symmetric organs with limited curvature, or the SAM, a dome-shaped structure exhibiting an anatomy of moderate complexity. However, not all plant organs fall into these simple morphogenetic categories. For example, strong curvature caused by developmentally regulated differential growth limits the usefulness of the implemented analytical strategies in iRoCS and 3DCellAtlas, particularly for indexing the axial position of a cell and determining its absolute distance to a reference. Yet, curvature represents a central element of the morphogenesis of plant organs with complex 3D shapes [28]. Many plant organs exhibit varying degrees of curvature, for example the apical hook of seedlings, leaves, or floral organs, such as sepals or petals. The ovule, the major female reproductive organ in higher plants, constitutes a particularly prominent example. Ovules are characterized by complex anatomy consisting of a central “trunk”, made up of several functionally distinct tissues stacked on top of each other, and by one or two laterally attached integuments, determinate planar tissues that eventually develop into the seed coat [29]. Moreover, most angiosperm ovules exhibit an extreme curvature due to asymmetric growth of the integuments [30].

Here, we used the ovule of *Arabidopsis thaliana* as a model to develop new generic strategies, implemented in MorphoGraphX, that allow the straightforward establishment of intrinsic coordinate systems in organs of simple or elaborate shapes. To this end we took advantage of a recently generated digital 3D reference atlas of ovule development in Arabidopsis [27]. We illustrate how such a coordinate system enables rapid annotation of cell identity and cell position in 3D and greatly facilitates the quantitative analysis of cellular features. We applied our strategies to an investigation of cell proliferation patterns in the ovule primordium and the cellular basis of differential integument growth. Finally, we demonstrate the broad applicability of the introduced concepts by providing proof-of-concept analysis for selected parameters in different plant organs of varying shape complexity.

## Results

### Curvature-related complications in the assignment of axial position

The analytical strategies for the positional annotation of individual cells relative to tissue organization depend on the structure under study. For example, the straightforward approaches employed in iRoCS and 3DCellAtlas involve cylinder coordinates and work very well for indexing axial position of cells relative to a reference in the root or hypocotyl, structures exhibiting limited curvature (Fig. 1A). However, asymmetric growth caused by differential cell proliferation and/or cell expansion can result in slanted or highly curved organs. In these instances, such approaches may lead to cells of the same indexed position having different absolute axial distances to a common reference (Fig. 1B). Thus, we devised a different strategy to minimize such axial distance errors when assigning 3D positional annotation to cells of organs that exhibit complex shapes. Our approach takes cues from central patterning events that frequently occur during early plant organogenesis, in particular the distinction of radial layers, the subdivision into anterior-posterior domains, and the establishment of a proximal-distal (axial) distance field (Fig. 1C,D). The proximal-distal distance of a cell relative to a user-defined reference is estimated by finding the shortest path through the cell centroids. Importantly, the search is confined to a given tissue layer and may not cross the anterior-posterior domain. The restriction to a tissue layer removes a large part of the axial distance error as the shortest path through the tissue layers cannot extend through interior tissues (Fig. 1C). On top of this restriction, prohibiting the shortest path from passing the anterior-posterior boundary further minimizes the error (Fig. 1D).

**Fig 1.**
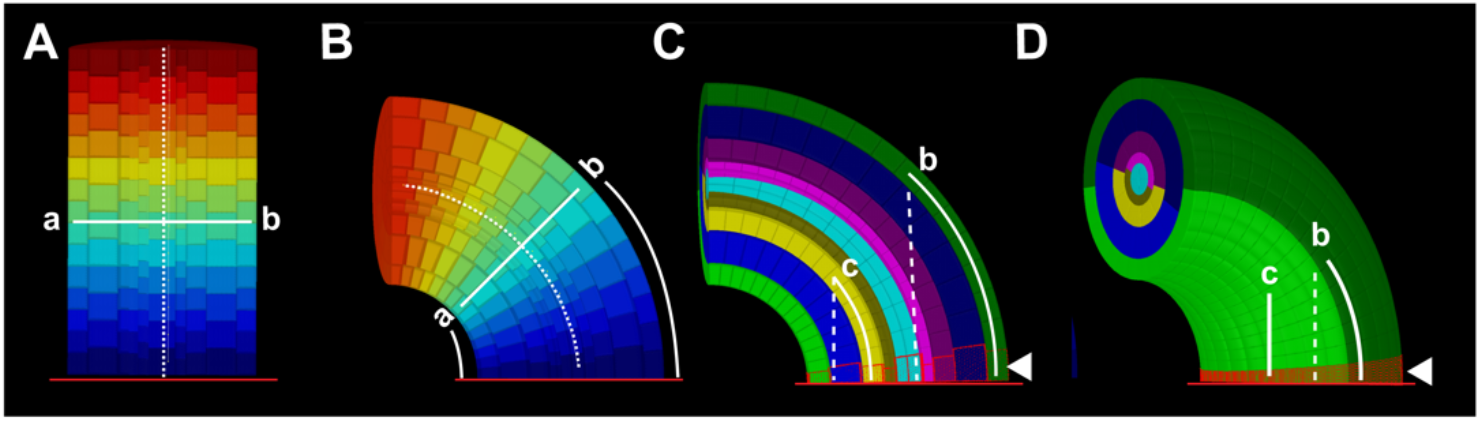
Axial cell distance determination in curved tissue. (A) Section through an artificial template of a tube-like and straight tissue consisting of multiple concentric cell layers. The heatmap indicates distance from the reference (red line at bottom). The dashed line outlines the central axis. Note that the two cells (a,b) at the same cell index position also show the same absolute axial distance to the origin (B) Same structure as in (A) but curved. Note that cells a and b differ in their axial distances to the reference. (C) Same structure as in (B). The separate cell layers are distinguished by their different colors. Two cells in different layers are highlighted (c, b). Dashed lines indicate shortest distances to the reference ignoring tissue layers. Solid lines mark the shortest distances to the reference that are restricted to tissue layers. Confining the shortest distance to a given layer reduces axial distance errors. The red line at the bottom highlights the reference. The arrowhead marks origin cells outlined in red. Origin cells exhibit a close distance in 3D to the reference (5-15 µm). (D) 3D representation of (C) revealing how the anterior- posterior boundary further minimizes the axial distance error for a cell in the posterior half of the structure.

### Development of the Arabidopsis ovule

The typical angiosperm ovule represents a prime example of an elaborately curved structure. Ovule development in *Arabidopsis thaliana* is well described [24,27,31,32] (Fig. 2A,B). During stage 1 the ovule emerges as a finger-like protrusion from the placenta (staging according to [27, 32]). Eventually, three elements can eventually be recognized along the trunk of the developing ovule: the nucellus at its tip, the chalaza in the center, and the funiculus at the bottom. The nucellus generates the large sub- epidermal megaspore mother cell (MMC) that will undergo meiosis during stage 2.

**Fig 2.**
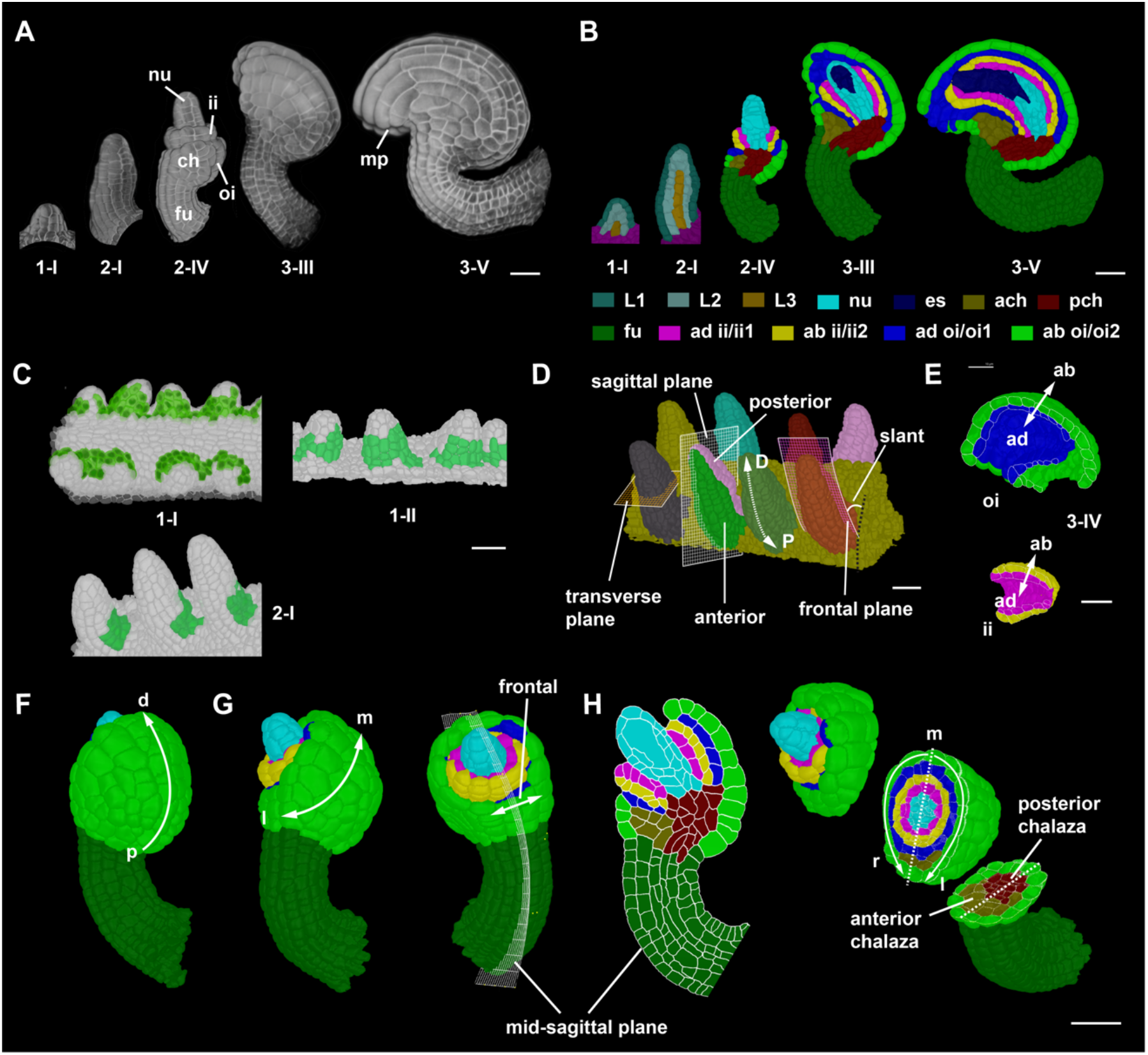
Developmental axes in ovule development. (A) 3D renderings of confocal z-stacks of SR2200-stained cell walls of wild-type ovules of the indicated stages. (B) Mid-sagittal sections through 3D digital ovules shown in (A). The different tissues are indicated. (C) 3D cell meshes highlighting the expression of the pKAN1::KAN1:2xGFP reporter in posterior epidermal cells from stage 1-I to 2-I in green. (D) 3D rendering of a placenta area carrying eight wild-type stage 2-I ovules. The anterior-posterior and proximal-distal axes are marked. The sagittal, transverse, and frontal planes are marked by grids on the primordia. Frontal plane separates the anterior and posterior halves of the organ. A Sagittal plane separates the left and right half of the organ. The dashed white arrow indicates the proximal-distal axis. Black dotted line indicates the placental surface to which the posterior side of the ovule is slanted with a small angle. (E) Outer and inner integument tissues extracted from the 3D mesh for visualizing the abaxial-adaxial polarity. Tilted view of mid-sagittal sections through the outer and inner integument, respectively, of a stage 3-IV 3D digital ovule. The arrows highlight the adaxial-abaxial axes of each integument. (F) Posterior view of a stage 2-V 3D digital ovule with the proximal-distal axis of the outer integument marked. (G) Side view (left) and anterior view (right) of the 3D digital ovule shown in (F). The medial-lateral axis of the outer integument and the frontal region are indicated. (H) A mid-sagittal section view (left) and a 3D clipped view (right) of the 3D digital ovule shown in (F) is depicted. It is oriented with the posterior side to the right. Tissue annotation as in (B). The 3D view allows the discrimination of the left-right sides of the 3D digital ovule. The dashed line indicates the medial line. Abbreviations: ab, abaxial; ad, adaxial; ach, anterior chalaza; ch, chalaza; ii, inner integument; es, embryo sac; fu, funiculus; mp, micropyle; nu, nucellus; oi, outer integument; pch, posterior chalaza; ml, medial-lateral; pd, proximal-distal; rl, right-left. Scale bars: 20 µm.

During stage 3 one of the meiotic products eventually develops into the haploid embryo sac or female gametophyte carrying the egg cell proper. The chalaza is characterized by two epidermally-derived integuments, lateral tissues that initiate from its flanks during stage 2. The outer integument represents a bilayered structure while the inner integument eventually consists of three cell layers. The two integuments grow around the nucellus but leave open a small cleft, the micropyle, through which the pollen tube can reach the embryo sac. The funiculus, a stalk-like structure, harbors the vascular strand and connects the ovule to the placenta. The mature Arabidopsis ovule features an elaborately curved shape. Curvature is caused in part by the integuments bending around the nucellus during stage 3 until their tips eventually locate next to the funiculus (anatropy). In addition, the funiculus forms a bend as well. Overall, the mature ovule exhibits a characteristic doubly- curved structure (Fig. 2A).

### Developmental axes of the ovule primordium

The Arabidopsis ovule primordium exhibits the typical radial organization into the L1, L2, and L3 cell layers [33, 34]. In addition, the distal nucellus, central chalaza, and proximal funiculus, represent three proximal-distal pattern elements along the trunk of the developing ovule. Gene expression patterns underlying the proximal-distal organization of the primordium are relatively well understood [27,35,36]. Importantly, the early primordium is not growing in a straight fashion but rapidly adopts a slant relative to the placenta surface, with the small angle of the slant facing the septum. It represents the first morphological sign of an anterior-posterior polarity (Fig. 1C,D) [27]. The presence of an anterior-posterior axis is further supported by the anterior expression of the class III HD-ZIP gene *PHABULOSA* in the early primordium [37]. To corroborate the establishment of an anterior-posterior axis in the stage 1-I ovule primordium we investigated the spatial signal distribution of pKAN1::KAN1:2xGFP, a reporter for *KANADI1* expression [38].

We observed detectable signal exclusively in the epidermis of the posterior ovule primordium (Fig. 2C). Interestingly, we did not detect expression in the tip of the primordium. By stage 2-I reporter signal appeared to be restricted to the posterior epidermis of the prospective funiculus.

In summary, the combined evidence strongly suggests the establishment of a radial, a proximal-distal, and an anterior-posterior axis in the ovule primordium. At the same time, the slant represents an early morphological manifestation of the ovule becoming a curved structure.

### Cell layer detection, individual organ separation, and anterior-posterior domain annotation in 3D

The annotation of cell position in 3D in the slanted ovule primordium required the application of the general principles outlined above. To be able to do so in a fast and robust manner we devised a new method for radial tissue labeling. Current pipelines, such as 3DCellAtlas and 3DCellAtlas Meristem, invoke a surface mesh as a central tool for cell and tissue annotation in the root, hypocotyl, and the SAM [10, 22]. However, establishing a surface mesh does not work well in situations where organs are in close contact with each other as is often the case for young ovule primordia attached to the placenta (Fig. 3A). The resulting individual surface meshes fail to outline the surfaces of the cells in contact and it is labour intensive to recreate a surface mesh in such instances.

**Fig 3.**
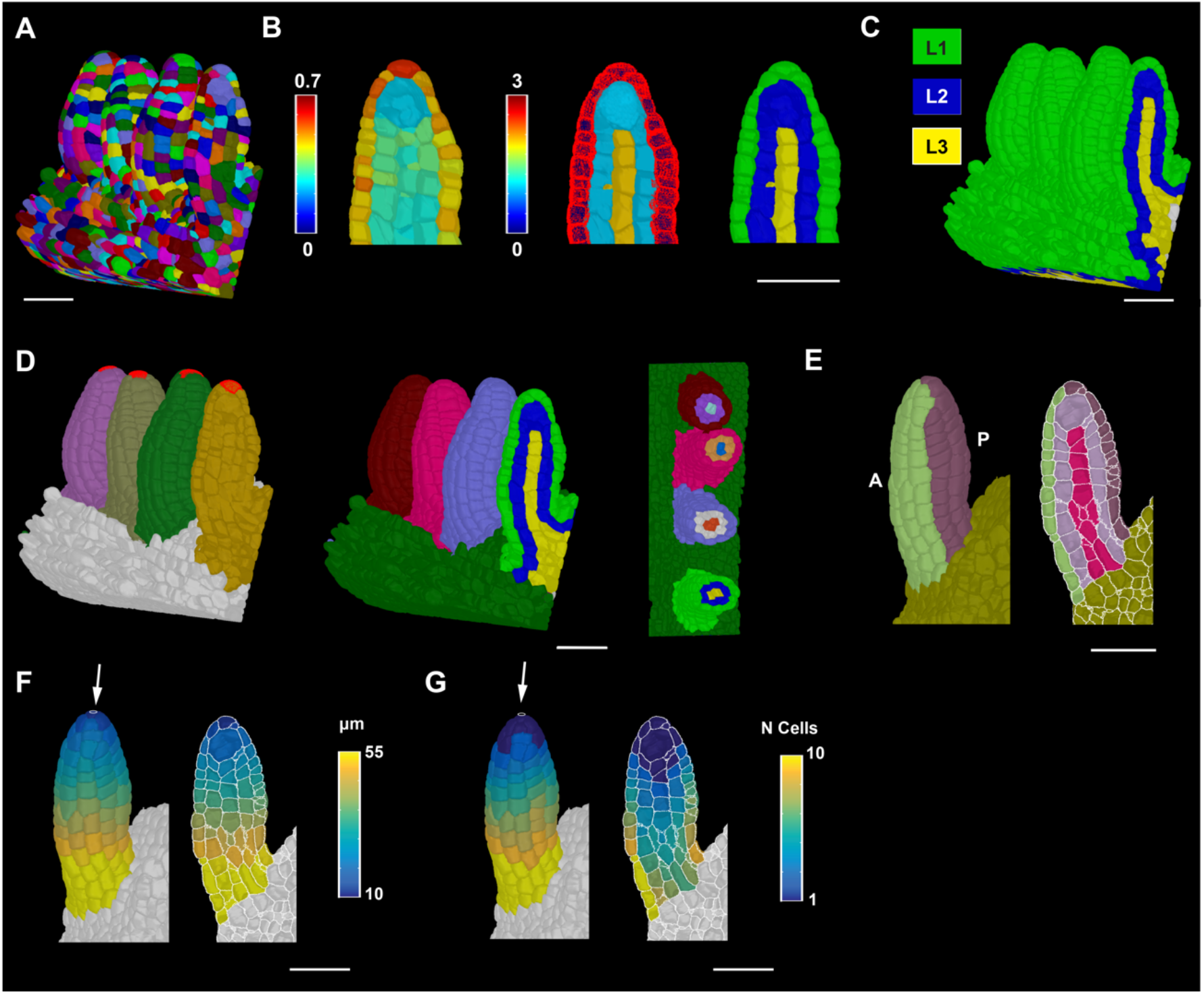
Ovule primordium tissue detection and coordinate system. (A) 3D segmented cell mesh view of a pistil fragment with four ovules of late stage 2-I. (B) Left panel: zoomed view of a sagittal section displaying the heatmap of outside wall area ratio. Threshold selection of outer surface cells based on the heatmap of outside wall area ratio. Center panel: heatmap indicates cell index, the number of cells an individual cell is separated from the selected outer surface cells marked in red. Right panel: heatmaps of cell distances (from center panel) were converted to integer values representing the tissue identity labels L1, L2 and L3. (C) Same method applied to the entire 3D mesh shown in (A). (D) Left panel: 3D Mesh view of the specimens shown in (A) with the distal most cell selected for organ separation. Center panel: colors on individual ovule primordia represent the results of organ separation after selecting the distal most cell and clustering the cell connectivity network. Result of the combination of L1, L2 and L3 label and organ separated labels annotated for the ovules shown in (A). Right panel: transverse section displaying the L1, L2, L3 labels for different ovules in different colors. (E) Anterior and posterior labels added to the tissue-annotated ovule primordia 3D cell meshes. Left panel: surface view. Right panel: mid-sagittal section. (F) Heatmap of distance coordinates from the point-like origin at the distal end of the organ (white arrow). Heat values indicate the distance in μm from individual cells centroid to the Bezier ring (indicated by white arrow) of the coordinates in a tissue restricted manner. Left panel: 3D view. Right panel: sagittal section view. (G) Heatmap of cell coordinates instead of distance coordinates as in (F). Heat values indicate how many cells apart is a cell of interest from the origin of the coordinates through tissue-restricted manner. Left panel: 3D view. Right panel: sagittal section view. Scale bars: 20 µm.

To address this problem, we developed a new strategy to perform automatic layer detection that groups cells into L1, L2, and L3 without a need for a surface mesh (Fig. 3A-C). In a first step L1 cells are clustered on the basis of a cell at the outer surface of the organ not being flanked by a neighboring cell at their outer surface. This feature is captured by defining the ratio of unshared wall area to shared wall area of individual cells (outside wall area ratio). Once L1 cells are clustered, L2 and L3 cells are found by their relative distances to the L1 cells. To this end a network of cell centroids is established and the shortest number of centroids a cell must cross to reach to the nearest L1 cell is determined. The corresponding result essentially reveals how many cells separate the cell of interest from the L1. The information can be directly used to cluster cells into L2 and L3 as L2 cells are direct neighbors of L1 cells and L3 cells are separated from the L1 by more than one cell. The strategy does not completely solve the issue when surface cells of neighboring organs are in full contact. However, the process works well when there is partial contact that still leaves behind a significant outside unshared cell wall area. This approach successfully annotated the radial tissue layers in ovule primordium or the shoot apical meristem (Fig. 3A-C, Fig S1).

Another problem relates to the separation of the multiple ovule primordia attached to the placenta into distinct units to allow ovule-specific analyses. We devised a method that takes advantage of the cell connectivity graph (Fig. 3D). From a selected cell at the distal end on each different ovule, distances to all other cells on the cell connectivity graph are computed. Cells are then assigned a label based on their nearest selected cell on that graph. A further parameter sets a maximum size of the ovules (in number of cells from the selected cells) to separate the ovules from their surrounding tissue. To facilitate downstream analysis, different labeling types, such as the cell layers and the ovule labels, are combined to create a unique label for each layer in every ovule.

In the last step, cells of anterior and posterior domains of about similar dimensions are obtained by manual selection (Fig. 3E). For the funiculus we also devised a semi-automatic method to distinguish the anterior and posterior domains (Fig. S2). In summary, the outlined approaches enable the generation of 3D digital ovule primordia of separate identities and near-perfect radial and anterior-posterior tissue annotation with minimal user input.

### Assignment of proximal-distal position to individual cells in 3D

To determine the proximal-distal (axial) position of an ovule primordium cell we generated a method that applies to 3D digital ovules for which radial cell layers and the anterior-posterior domains have already been annotated. The proximal-distal position of each cell is calculated, either in terms of cell index or absolute or relative distance to a reference, in a two-step procedure (Fig. 3F,G). In the first step, a Bezier ring (a mathematically defined curved line) will serve as reference for the proximal-distal distance field and is placed at one end of the tissue. In the case of the cone-shaped ovule primordium a small, near point-size Bezier ring is positioned at the distal tip of the primordium. Positioning the Bezier ring at the distal tip correlates with a biologically relevant maximum of the phytohormone auxin at the tip as inferred from the expression of the auxin response reporter pDR5::GFP [39], the spatial signal of the auxin sensor R2D2 [40, 41], and the finding that polar auxin transport mediated by *PINFORMED1* (*PIN1*) is required for ovule primordium formation [42, 43]. In case of the mature funiculus that is close in shape to a curved cylinder, a larger Bezier ring is placed at its proximal end. Origin cells are then defined by their close distance in 3D (usually 5-15 µm) to the user-specified Bezier ring (Fig. 1C,D). They act as seeds for the distance coordinates of the remaining cells of the tissue that are obtained by searching for the shortest path through the cell centroids to the centroid of an origin cell. The search is restricted to a radial layer and by the anterior-posterior boundary. It should be noted that with this approach small axial distance errors still occur within the anterior or posterior domains depending on the number of laterally arranged cell files within these areas. The remaining errors are typically in the range of a few microns, but can be eliminated when taking individual cell files into account. This is possible within the software, however, the procedure involves cumbersome manual annotation of all cell files for each cell layer. Moreover, the gain of resolution is minimal.

### A coordinate system for integuments

The two integuments undergo complex morphogenesis with the inner integument eventually resembling a curved cylindrical shell and the outer integument developing into a curved hood-like shell (Fig. 2A,B,F). Both integuments are characterized by their own intrinsic developmental axes (Fig. 2E-H). A distinct adaxial-abaxial (dorso-ventral) axis is prominent as the individual cell layers differ in cellular morphology and gene expression patterns [32,37,44–46] (Fig. 2E). Both integuments also feature their own proximal-distal axes (Fig. 2A,F). Related to its hood-like shape the outer integument flanks a frontal section and features a medial-lateral axis (Fig. 2G). As a rule, we position the ovule with the anterior domain and the micropyle pointing to the left and the proximal end of the funiculus pointing towards the bottom right. Based on this arrangement we define the left and right lateral sides of the medial-lateral axis (Fig. 2H).

We successfully applied to the integuments some of the same formal strategies as described for the primordium or funiculus. In an initial step, the integumentary adaxial- abaxial cell layers are labelled manually. In a subsequent step, medial-lateral coordinates of all cells of a given integument layer are established relative to a file of posterior midline cells (Fig. 4A). Cell distance is computed in terms of how many cells separate a given cell from the midline (Fig. 4B). Cells along the medial-lateral axis can be grouped further into median and lateral subdomains that occupy about half the width of an integument layer. For example, for the outer layer of the outer integument we grouped cells that are located three cells to the left or right of the posterior midline cells into the median domain. The remaining cells are classified as lateral cells (Fig. 4C). In the following step, proximal-distal distance coordinates are assigned for all integument cells. A Bezier ring is first placed at the proximal end of the inner side of the outer integument (next to its inner layer) facing the outer layer of the inner integument (Fig. 4D). The circular origin is in the same plane as the ring-shaped expression pattern of the pCUC3::CUC3:CFP reporter which marks the proximal base of the two innermost layers of the inner integument, respectively (Fig. 4E). Members of the CUC gene family are generally required for primordium initiation and organ boundary formation [43,47–52]. Origin cells are then defined by their close user-specified distance to the Bezier ring in 3D (about 5-15 µm). As a direct result of the placement of the Bezier ring cells of the outer layer of the outer integument that are in direct contact with the subepidermal proximal chalaza are assigned a negative value for the proximal-distal position (Fig. 4F). This feature can be used to separately cluster and analyze those cells. Finally, proximal-distal distance coordinates of the integument cells are obtained by searching for the shortest path through the cell centroids to the centroid of an origin cell (Fig. 4F,G). The search is again restricted to a given tissue layer and may not cross the medial-lateral boundary. Taken together, the procedure assigns medial-lateral and proximal-distal positions for all cells of the integuments.

**Fig 4.**
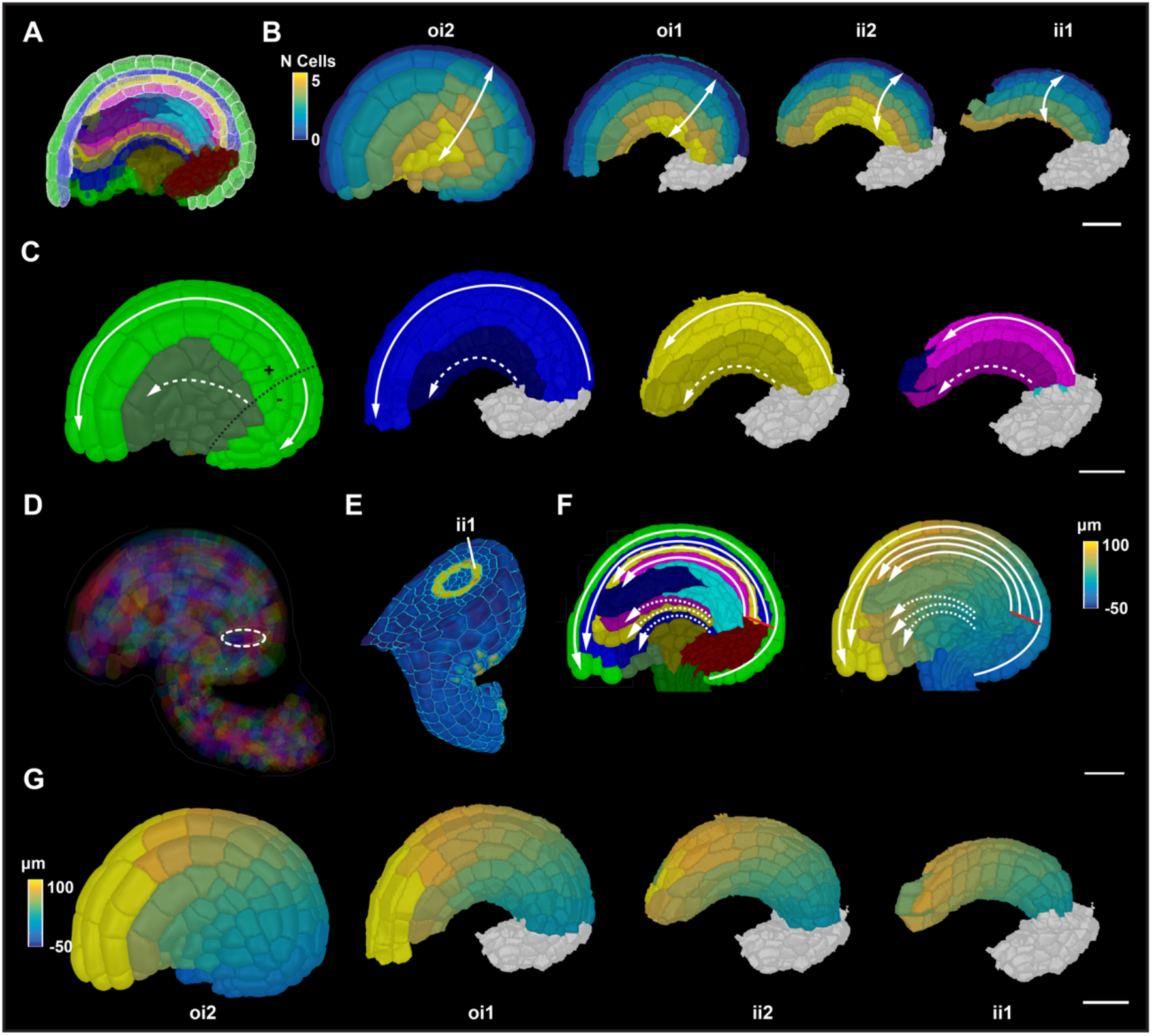
Integument coordinate system. (A) Mid-sagittal section highlighting the selected medial cells on the posterior side of the four layers of integument tissues for medial-lateral coordinate annotation. Colors represent tissue annotations similar to Fig. 1B. (B) Heatmap of medial-lateral cell coordinates. Heat values indicate the lateral position in terms of the number of cells from the median file of cells. Different integument tissues are extracted from the 3D mesh to display the medial-lateral coordinates at their tissue surface. (C) 3D surface view of integument tissues similar to (B). Medial and lateral cells are distinguished. Solid white line represents the tissue restricted coordinate direction along the medial group of cells. White dashed line represents the tissue restricted coordinate direction along the lateral group of cells. Black dotted line on oi2 represents the coordinate origin projected on the surface which separates the proximal oi2 cells with negative coordinate values (D) Semi-transparent view of a mature 3D ovule displaying the coordinate origin as a ring inside the organ. (E) 3D clipping view of a transverse section of an ovule highlighting the ring-like expression of the pCUC3::CUC3:CFP reporter in yellow. (F) Left panel: sagittal section of a mature ovule displaying the coordinate directions of the medial and lateral group of cells in solid and white lines, respectively. Solid red line indicates the origin of the coordinate system. Right panel: Sagittal section displaying the heatmap of distance coordinates. Solid red line indicates the origin of the coordinate system. (G) 3D surface view of integument tissues similar to (A) displaying the distance coordinates at the surface of internal tissues. Scale bars: 20 µm.

### Differential distribution of cellular growth patterns during early ovule development

To provide proof of concept for the usefulness of our computational tools in the quantitative analysis of cellular patterns in a 3D context we assessed spatial growth patterns in selected aspects of ovule development. To this end we made use of a previously published dataset of wild-type 3D digital ovules of the Col-0 accession [27]. We first focused on primordium outgrowth. It was previously shown that ovule primordia grow in a continuous fashion based on an analysis of the total number of cells and the increase of organ volume from stages 1-I to 2-I [24, 27]. However, it remained unclear how cell numbers and cell volumes of the radial layers compare to each other. In addition, it was not known if mitoses are randomly distributed along the proximal-distal axis or if they preferentially occur in specific domains. To address these questions we analyzed 52 3D digital wild-type ovule primordia that encompassed stages 1-I to 2-I (Fig. 5A,B).

**Fig 5.**
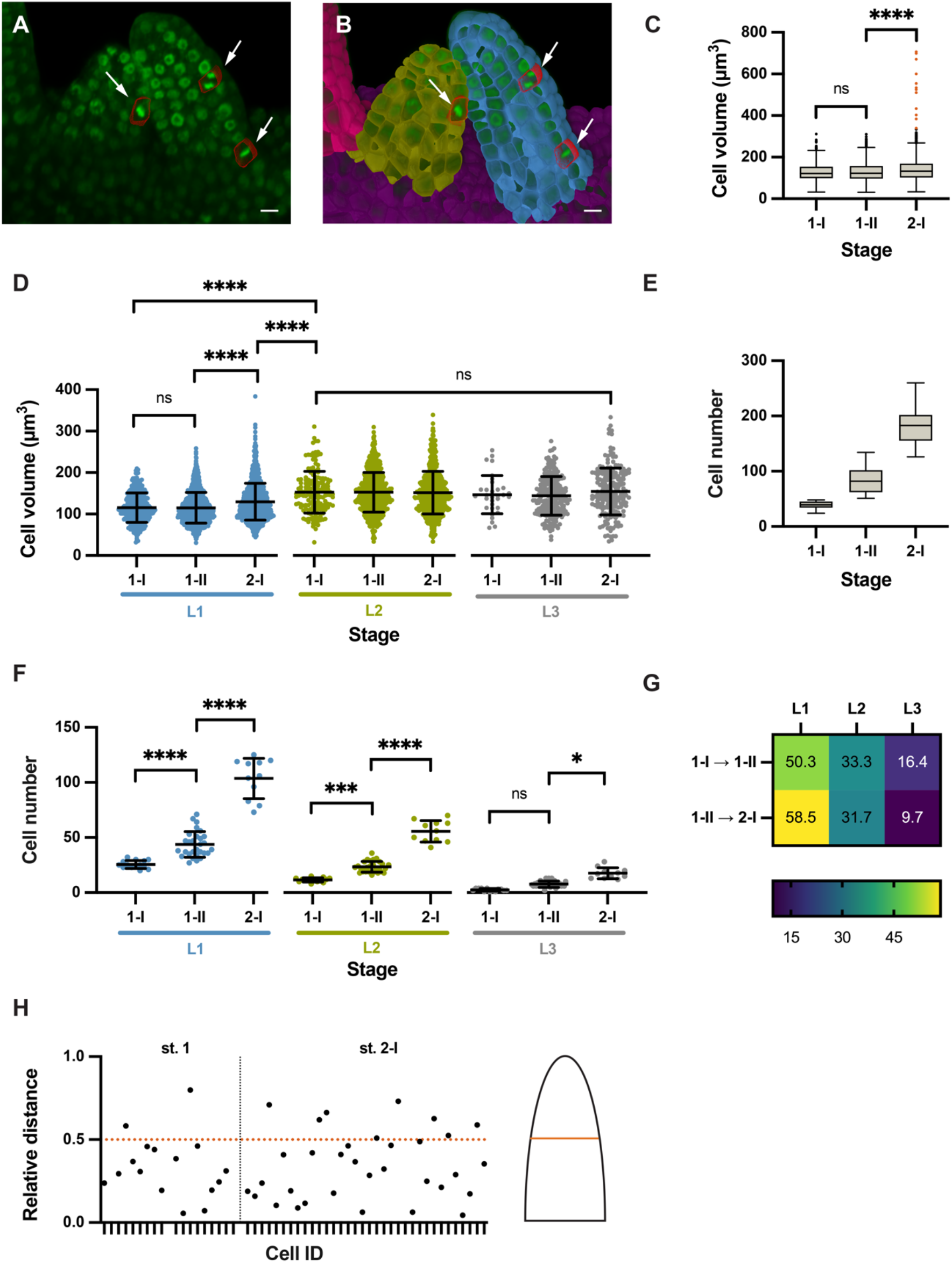
Cellular growth patterns in the ovule primordium. Different stages of wild-type (Col-0) ovule primordia are analyzed. (A) 3D frontal plane view of ovule primordia displaying the To-PRO-3 nuclear stain. Cells undergoing mitosis are outlined in red and marked by arrows. (B) Same section in (A) with an overlay of 3D labelled meshes. Arrows indicate mitotic figures (C) Comparison of cell volumes. Box and whiskers plots are shown. Whiskers extend to 1.5x the interquartile range from the 25 or 75 percentile, respectively (Tukey). (D) Comparison of cell volume between radial layers of different stages. (E) Comparison of cell numbers. Box and whiskers plots are shown. Whiskers extend to 1.5x the interquartile range from the 25 or 75 percentiles, respectively (Tukey). (F) Comparison of cell numbers between radial layers of different stages. (G) Heatmap of percentage increase in number of cells per tissue compared to overall increase in number of cells. (H) Plot showing the relative distance along the proximal-distal axis of mitotic cells in a primordium. Relative distance was calculated by the following formula: reldist = 1 - distance coordinate/mean length. The stages are indicated. Statistics in (D, F): data are mean ± SD; significances: ****, p < 0.0001; ***, p < 0.0004; *, p < 0.02. Ordinary one-way ANOVA followed by Tukey’s multiple comparison test. Scale bars: 20 µm.

We initially undertook a comparison of cell volumes between stages (Fig.5C) and radial layers (Fig. 5D) (Table 1). In this dataset the L2-derived MMCs at stage2-I feature an average cell volume of 543.3 µm^3^ ± 120.6 µm^3^ (mean ± SD) with a minimal volume of 335 µm^3^ [27]. Thus, the volume of the MMCs is beyond the largest cell volumes observed for other cells (Fig. 5C). We therefore eliminated the MMCs from this analysis to eliminate skewing of the results due to their out-of-range size. We observed that the average volume of L1 cells slightly increased during development while the average volume of L2/L3 cells stayed about constant (Fig. 5D) (Table 1). The results further indicated that with the exception of the MMCs the L2 and L3 feature cells of about similar cell volumes while the L1 is composed of smaller cells.

**Table 1.** Layer-specific cellular growth characteristics in the ovule primordium.

| Cell layer | Stage<sup>a</sup> |  |  |  |  |  |
| --- | --- | --- | --- | --- | --- | --- |
|  | I-I |  | 1-II |  | 2-I |  |
|  | Cell number | Cell volume<sup>b</sup> | Cell number | Cell volume<sup>b</sup> | Cell number | Cell volume<sup>b</sup> |
| L1 | 25.6 ± 3.7 | 115.6 ± 35.7 | 43.8 ± 11.6 | 115.2 ± 36.8 | 103.6 ± 18.4 | 129.9 ± 44.5 |
| L2 | 11.6 ± 1.9 | 152.8 ± 50.3 | 23.4 ± 4.9 | 152.7 ± 47.6 | 55.6 ± 9.7 | 158.5 ± 74.2 |
| L3 | 2.5 ± 1.2 | 146.8 ± 45.9 | 7.7 ± 2.8 | 144.1 ± 46.6 | 17.6 ± 5.0 | 154.7 ± 56.5 |

<sup>a</sup>Number of 3D digital ovules scored: 14 (stage 1-I), 28 (stage 1-II), 11 (stage 2-I).
<sup>b</sup>Volumes are given in $\mu\text{m}^3$ .
Values represent mean ± SD

We then compared cell numbers between stages (Fig. 5E) and radial layers (Fig. 5F) (Table 1). Overall, we observed a steady increase in cell numbers during primordium outgrowth (Fig. 5E). The L1 covers a larger surface of the primordium than the L2 or L3. Considering this aspect and given the smaller cell volume of L1 cells in comparison to L2/L3 cells, we hypothesized that the L1 consists of more cells than the L2 and L3. This assumption was supported by layer-specific cell counts (Fig. 5F) (Table 1). We also determined that the L1 showed the largest percentage increase in cell numbers while the L3 showed the least percentage increase in cell numbers (Fig. 5G).

Next, we assessed the spatial distribution of mitoses. To this end we manually labelled cells that exhibit mitotic figures (Fig. 5A,B). We identified a total of 52 mitotic cells in our dataset comprising 52 3D digital ovule primordia. We first asked if there were differences in the number of mitoses between the cell layers. We found 33 mitotic cells in the L1, 18 in the L2, and 1 in the L3. This result is in line with the observed differences in cell numbers between the three layers (Fig. 5F). We then investigated if there was a difference between the number of mitotic cells in the anterior and posterior L1. We found 23 and 10 mitoses in the anterior and posterior L1, respectively, indicating that more cell divisions occur in the anterior domain. Finally, we analysed the proximal-distal distribution of mitoses. We found that about 80 to 85 percent of scored mitotic cells were located in the proximal half of the developing primordium (Fig. 5H).

Taken together, our data indicate that primordium outgrowth is preferentially driven by an increase in cell number, not cell volume. In addition, they suggest unequal spatial distribution of mitoses between cell layers and along the anterior-posterior and proximal-distal axes. A higher number of mitoses in the anterior domain might explain primordium slanting. The data further indicate that a cell proliferation zone located in the bottom half of the developing primordium contributes significantly to its outgrowth.

### Funiculus curvature correlates with differences in cell number and cell volume along the anterior-posterior and proximal-distal axes

To obtain insight into the cellular processes underlying funiculus curvature we performed an initial analysis of its cellular characteristics using 14 3D digital ovules of stage 3-IV (Fig. 6A-E). By this stage growth of the funiculus has ceased [27]. We focused on the L1 and L2 layers. To investigate cellular features of the L2 we digitally removed the L1 from the 3D digital funiculi. First, we assessed the proximal-distal distance and cell number along the L1 midlines of the anterior and posterior domains, respectively (Fig. 6A-E). We observed that the anterior midline was longer and characterized by a higher number of cells in comparison to the posterior midline. To directly compare volumes of anterior and posterior cells we converted the cells’ coordinates into relative proximal-distal positions [53]. There we noticed a gradient in the volume of anterior L1 and L2 cells with the distal-most cells featuring nearly 1.5 to 2 times the volume of cells located at the proximal end (Fig. 6D,E). We did not detect a noticeable volume increase in the posterior cells.

**Fig 6.**
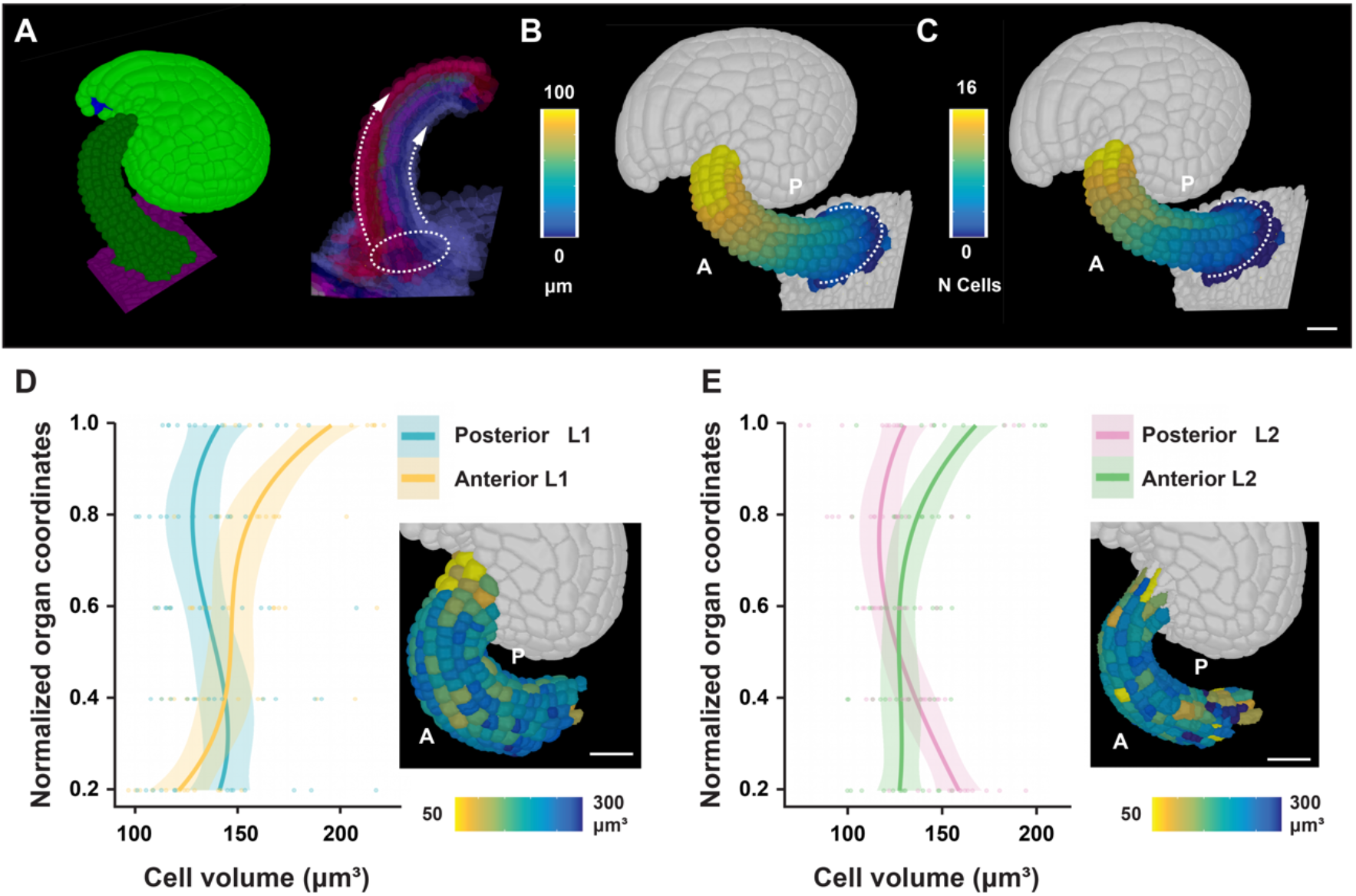
Cellular features of funiculus curvature. Wild-type (Col-0) ovules of stage 3-IV are analyzed. (A) Left panel: Tilted side-view of a 3D cell mesh. Right panel: Semi- transparent 3D mesh view of funiculus extracted from the 3D mesh of the organ. The Bezier ring serving as origin is placed at the proximal base of the funiculus. The dashed arrow lines indicate the coordinate direction along the anterior and posterior midlines. (B,C) Same specimen as in (A). The anterior and posterior sides are marked. (B) Heatmap of cell distances along the proximal-distal axis of the funiculus. (C) Heatmap of cell numbers along the proximal-distal axis of the funiculus. White dotted line indicates the coordinate origin as a ring. (D) Graph depicting cell volumes of anterior and posterior L1 cells in relation to the normalized proximal-distal position. The inset in the bottom right corner shows a heat map of cell volume in the L1 of the funiculus. 14 3D digital ovules were analyzed. Regression curves are cubic polynomials with 95% confidence intervals (shaded regions). Number of cells: 1568 ≤ n ≤ 1768. (E) Similar graph as in (D) revealing cell volumes of anterior and posterior L2 cells. Number of cells: 1175 ≤ n ≤ 1352. Scale bars: 20 µm.

In summary, our data suggest that a combination of differential cell proliferation along the anterior-posterior axis and unequal cell growth along the proximal-distal axis of the anterior domain contributes to the curvature of the funiculus.

### Proximal-distal growth gradient in Arabidopsis integuments

Genetic data indicated that asymmetric growth of the outer integument contributes significantly to the anatropous shape of the ovule [27,54,55]. However, the 3D architecture of integument cells in relation to their position within the tissue remained unknown. Thus, we undertook a first analysis of the 3D geometry of integument cells in 3D digital ovules of stage 3-IV. At this stage curvature is underway but not yet completed. We used 3DCoordX to measure cell volumes of the outer layer of the outer integument along the proximal-distal axis. We found a gradient in cell volume along this axis. We observed that proximal cells exhibited small cell volumes while, with the exception of small cells at the tip of the integument, distal cells were characterized by larger sizes (Fig. 7A). Next, we expanded the 3D cellular analysis to all cells of the integuments. In a typical stage 3-IV 3D digital ovule we found a proximal-distal gradient of cell volumes in cells of the other layers, but at a smaller scale compared to the outer layer of the outer integument (Fig. 7A-C). We then compared cell length to cell position along the proximal-distal axis in the medial domain of the inner layer of the outer integument across five different specimens. We found that cell length increased along the proximal-distal axis (Fig. 7D).

**Fig 7.**
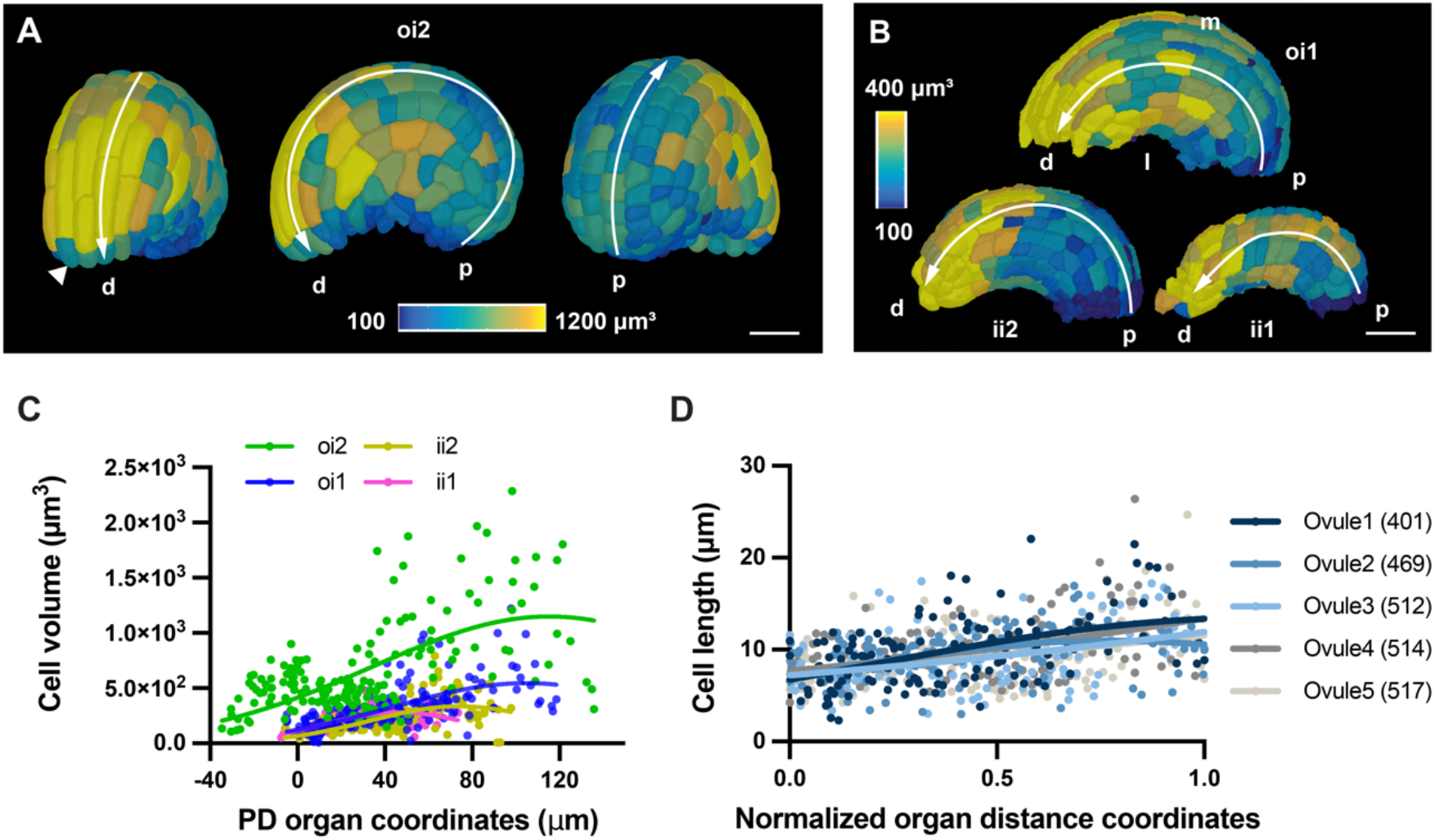
3D geometry of integument cells. Wild-type (Col-0) ovules of stage 3-IV are analyzed. (A) Heatmap of oi2 cell volumes. Panels depict a tilted frontal view (left), side view (center), and tilted back view (right). The proximal-distal axis is indicated. The arrowhead marks a small tip cell. (B) Side views of the same specimen as in (A) showing the 3D surface view of internal tissues. Heatmaps of the oi1, ii2, and ii1 layers, respectively. (C) Graph showing cell volume in relation to proximal-distal distance for the four integument layers of a representative ovule. The respective nonlinear Gaussian regression curves are indicated. 95 ≤ n ≤ 172. (D) Cell lengths in relation to normalized proximal-distal distance. Cells in the median oi1 layer of five ovules were analyzed. The respective nonlinear Gaussian regression curves are indicated. 126 ≤ n ≤ 133. Note the increase in cell length towards the distal end. Abbreviations: d, distal; ii1, inner layer of inner integument; ii2, outer layer of inner integument; l, lateral; m, medial; oi1, inner layer of outer integument; oi2, outer layer of outer integument; p, proximal.Scale bars: (A,B) 20 µm.

Taken together, our data suggest that preferential cell elongation along the proximal-distal axis may be an important factor underlying differential growth of the outer integument and ovule curvature.

### Application to other plant organs

Finally, we explored if our approach for an organ-intrinsic coordinate system was useful beyond the Arabidopsis ovule and could be of value to provide spatial context to cellular growth patterns in different organs of various plant species. To this end we investigated 3D digital plant organs of diverse morphological complexity. We first inspected the archegonium of the liverwort genetic model system *Marchantia polymorpha*. The archegonium is an organ of simple morphology consisting of two main tissues: the spherical venter harboring the egg cell and the elongated neck through which the sperm cell reaches the egg cell [56]. We generated two 3D digital archegonia: a younger specimen A and an older specimen B. Both archegonia were imaged, 3D segmented, and cells of the neck, neck canal, venter, and venter canal were identified and labelled manually. We removed the egg and canal cells from our analysis and focused on the venters and the necks of the two 3D digital archegonia (Fig. 8A,B). A first inspection already revealed differences between the two specimens. We observed that the venter of specimen A possessed only one cell layer with no obvious signs of periclinal cell division. By contrast, we found that a scattered pattern of periclinal cell divisions had occurred in the venter of specimen B associated with the formation of a second cell layer (Fig. 8B) [56]. This observation indicated temporal and spatial asynchrony in the control of these periclinal cell divisions.

**Fig 8.**
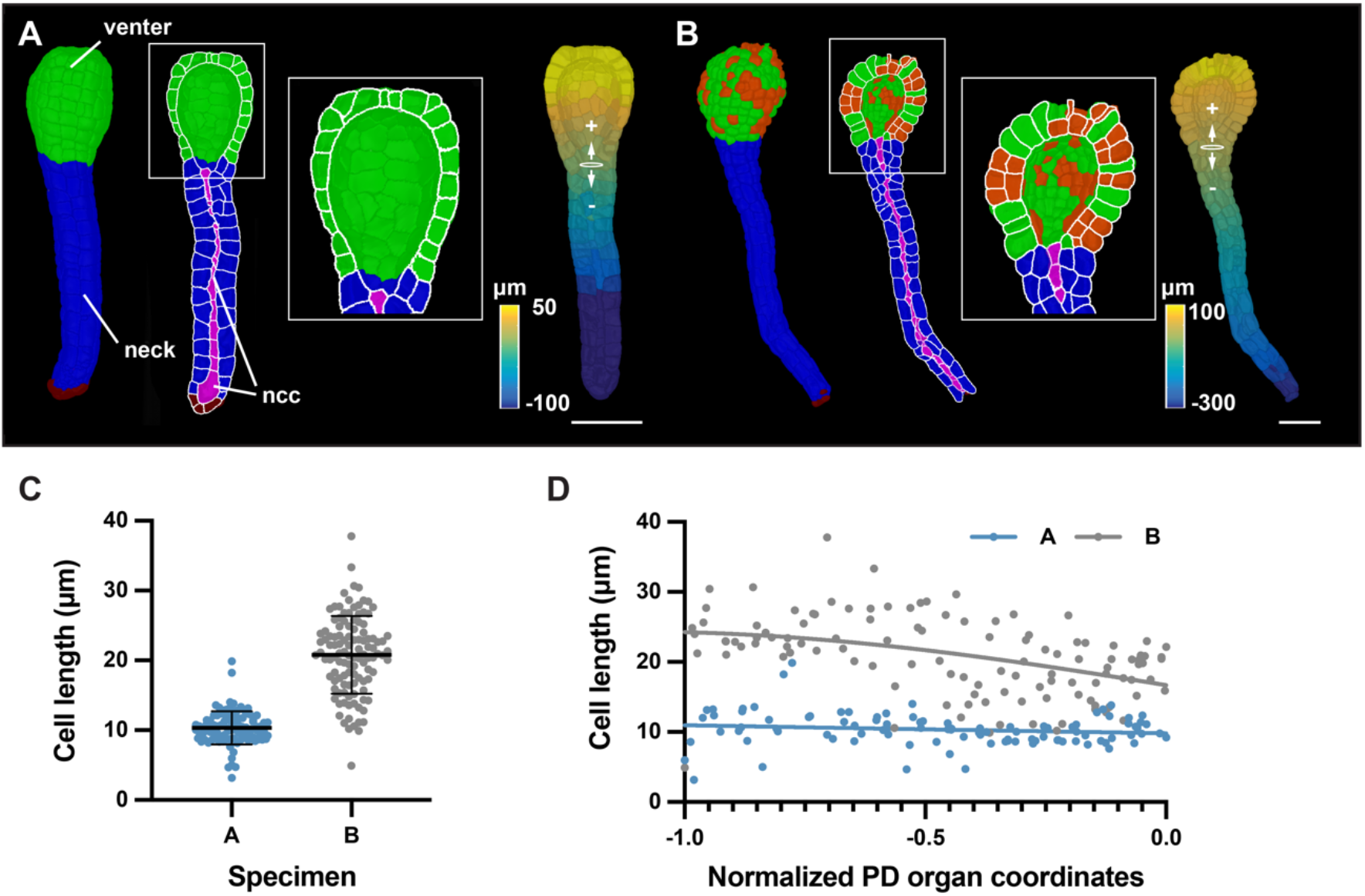
Cellular analysis of 3D digital archegonia from Marchantia. (A,B) 3D cell meshes of two different-stage archegonia are depicted. (A) Specimen A. The venter and neck are indicated. Left: 3D view. Center: An about mid-sagittal section. Right: Distance values along the central axis. The Bezier ring at the venter-neck boundary, which serves as the origin of the coordinate system, is indicated. (B) Specimen B. Identical arrangement as in (A). Highlighted white box represents the zoom view of the venter cells having undergone periclinal cell divisions marked in red. (C) Graph depicting the cell lengths of all neck cells of specimens A and B. The mean ± SD is indicated. Specimen A: n = 98. Specimen B: n = 114. (D) Graph showing cell length of individual neck cells of the two specimens shown in (C) in relation to their normalized position along the long axis of the neck. The tip of the neck is oriented to the left (0.0 corresponds to the ring position, the tip of the neck is at position -1.0, compare with Fig. 8B). Abbreviations: ncc, neck canal cells. Scale bars: (A,B) 50 µm.

A monolayer of cells formed the necks of both specimens. We implemented an organ coordinate system to enable a spatial analysis of some basic cellular parameters along the long axes of the two specimens. We placed a Bezier ring at the boundary between the venter and neck cells (Fig. 8A,B). The placement of the ring resulted in the assignment of positive and negative organ coordinate values for the venter and neck cells, respectively. We then assessed the basis of the differences in neck lengths between the two specimens. Measuring neck length along the main central axis revealed that the neck of specimen B was about 2 times longer than specimen A (329 µm versus 160 µm). We then asked if the length disparity between the necks of the two specimens was due to a difference in cell numbers and/or cell elongation. We determined 98 and 114 neck cells for specimens A and B, respectively, indicating a minor difference in neck cell numbers. Next, we quantified cell length along the central organ axis for all neck cells. We observed that the neck cells of specimen A had an average length of 10.3 µm ± 2.4 µm and exhibited a relatively uniform cell length (Fig. 8C). Neck cells of specimen B showed a more heterogeneous distribution of cell length and were noticeably more elongated with an average cell length of 20.8 ± 5.6 µm. Moreover, cell elongation increased towards the tip of the neck in specimen B while no such increase was observed for specimen A (Fig. 8D).

The results indicated that enhanced cell elongation along the central axis of the neck was mainly associated with the increase in neck length in specimen B in comparison to specimen A.

Finally, we turned our attention to a plant organ of complex 3D morphology. The cup- shaped trap of the aquatic carnivorous plant *Utricularia gibba* represents a highly curved 3D leaf form [57, 58]. Quantitative growth analysis at the tissue level combined with computer modeling indicated that the complex shape transformations occurring during trap development are associated with differential rates and orientations of growth [59, 60]. However, a quantitative analysis of 3D cellular parameters had not been performed. To obtain first insight into the cellular basis of the growth patterns shaping the Utricularia trap we generated a 3D digital representation with cellular resolution of an intermediate- stage trap collected 6 days after initiation [59]. By this stage an invagination in the near- spherical young trap had occurred, followed by the formation of further folds and tissue broadening, and resulting in the emergence of the interior trap door and threshold (Fig. 9A-C). We manually labelled the various tissues, including the abaxial and adaxial cells of the wall, the threshold, and the combined trap door/palate domain, and distinguished between medial and lateral domains of the adaxial and abaxial wall, respectively. To define an origin of the distance coordinate system we placed an ellipsoid Bezier ring at the boundary between the base of the threshold and the wall of the trap (Fig. 9D). We then asked if there were position-related differences in cell volume in the epidermal layers of the wall and threshold by analyzing epidermal cells located along the respective midlines of the tissues (Fig. 9E-G). We observed that cell volume varied along the measured distances. For example, we noticed a sudden increase in cell volume in an interval from 240 µm to 320 µm for cells of the abaxial wall (Fig. 9F). This region precedes a prominent kink in the abaxial wall (Fig. 9E). By contrast, cell volumes of the adaxial wall dropped towards the end. The decrease in cell volume was likely associated with the tapering of the adaxial wall that could be observed in this area. Volumes of threshold cells positioned within a range of 80 to about 150 µm from the origin showed relatively small volumes in comparison to the cells flanking this interval (Fig. 9G). The 80-150 µm zone corresponded to a section of the threshold which was only moderately curved and provided a large surface exposed to the lumen of the trap (Fig. 9E). Taken together, the data revealed spatial differences in cell volume for all three examined tissues of this specimen and support the notion that differential cell growth contributes to the morphogenesis of the Utricularia trap.

**Fig 9.**
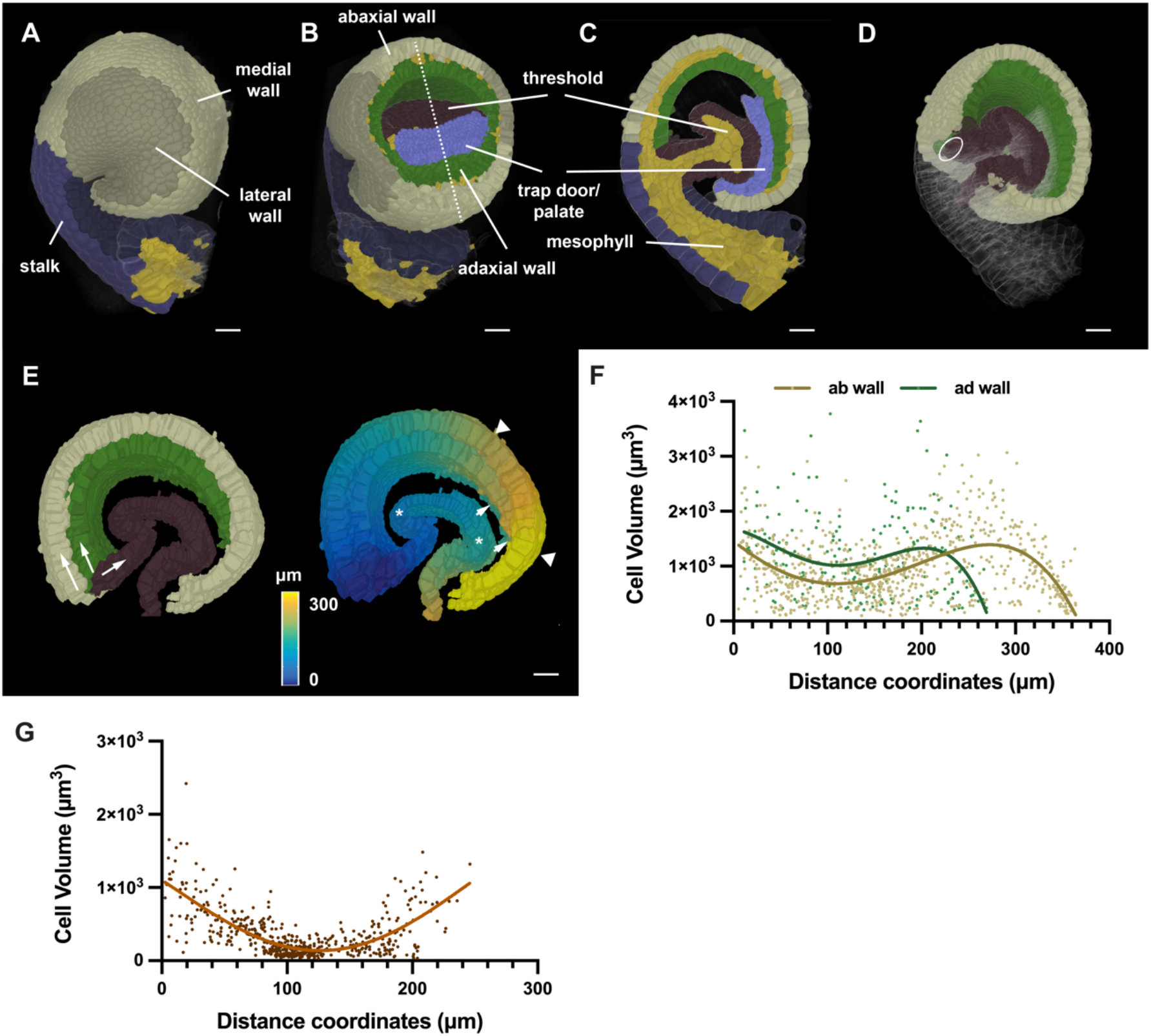
3D digital Utricularia trap. A specimen 6 days after initiation is shown. (A) Side view of the 3D cell mesh with annotation of various tissues. (B) Tilted view of (A) with part of the wall removed by a tangential clipping plane. The dashed line indicates the mid-sagittal section shown in (C). (C) Mid-sagittal section. (D) Slanted 3D view of (A) with half of the trap cut off at the mid-sagittal plane shown in (B). The trap door and palate domain were removed. The position of the Bezier ellipsoid is indicated. (E) Left panel: arrows indicate the direction of the distance coordinates through the epidermis of the abaxial and adaxial tissue of the wall and the threshold, respectively. Right panel: Heat map indicating distances. Wall and threshold are treated separately. Triangles mark the 240-320 µm interval of the abaxial wall. Arrows highlight the tapering end of the adaxial wall. Asterisks indicate the 80-150 µm interval of the threshold. (F) Graph displaying cell volume of epidermal trap cells in relation to their position. Values for the abaxial and adaxial wall are superimposed. The respective nonlinear regression curves fitting a fourth order polynomial function are indicated. ab wall: n = 786, ad wall: n = 231. (G). Graph displaying cell volume of threshold cells in relation to their position. The line marks a nonlinear regression curve. n = 533. Scale bars: 20 µm.

## Discussion

The generation of biological form can be explained in terms of growth oriented relative to an organ-centric coordinate system [7,28,61]. To understand tissue morphogenesis it is therefore essential to provide spatial context to the quantitative analysis of the molecular and cellular networks that underlie the development of tissues and organs. It requires robust methods that enable the objective assignment of position to individual cells in a rapid and reliable manner. Here, we developed 3DCoordX, a collection of computational tools that enable the assignment of organ-centric coordinate systems to several different plant organs with complex shapes that were not accessible with previous methods. By applying mathematically defined criteria for the annotation of cell position in 3D the tools largely eliminate user-derived ambiguities in the cell annotation process. 3DCoordX is implemented as an add-on to the open-source software MorphoGraphX [11, 14] (preprint). A detailed user guide can be found in the supplement. 3DCoordX enables quantitative analyses of cellular features in their spatial context, in a rapid manner, and on a large scale.

We took advantage of the recently made available 3D digital reference atlas of the Arabidopsis ovule to develop generic conceptual and computational approaches that enable the assignment of 3D coordinate systems to organs of simple as well as complex curved morphology. Previous efforts, such as iRoCS [9] and 3DCellAtlas [10], relied on externally imposed coordinate systems. The design of the strategies for the annotation of cell position presented here is guided by intrinsic patterning processes and assign cell distance in relation to organ-centric developmental axes. Such tissue polarity axes are thought to play a central role in the spatial control of growth [28,62,63]. To accommodate the particular architecture of the ovule we devised two 3D coordinate systems, one for the main “trunk”, the central proximal-distal axis consisting of the nucellus, chalaza and funiculus, and one for the integuments. Both coordinate systems are based on similar general principles. First, we distinguish between individual cell layers as in the L1 to L3 layers of the primordium or the adaxial-abaxial (dorso-ventral) cell layers of the two integuments. Importantly, with the concept of “outside wall area ratio” 3DCoordX embodies a new strategy for the classification of the radial layers. It does not rely on a surface mesh for classification and thus is more versatile than other published methods.

Second, the cell layers then become subdivided into two domains: the anterior-posterior domains of the trunk and the medial-lateral domains of the integument cell layers. Third, subsequent assignment of a proximal-distal distance value to each cell is constricted by these two prepatterns. For the placement of reference Bezier rings we took cues from the localization of important developmental regulators, such as the presence of an auxin maximum at the distal tip of the ovule primordium or the *CUC3* expression in the chalaza [39–41,52]. As a result of the approach each cell is annotated in 3D with respect to the radial and proximal-distal dimensions as well as to either an anterior-posterior or medial-lateral axis. With the help of 3DCoordX we readily discovered previously unidentified cellular growth patterns in the primordium and integuments. For example, our data indicate a basal cell proliferation zone in the ovule primordium and suggest that preferential cell proliferation in the anterior domain is important for primordium slanting. Moreover, we obtained evidence that the increase in cell volume along the proximal-distal axis of the outer integument, a tissue without radial symmetry, is mainly explained by an increase in cell length.

Importantly, our work revealed that the respective principles can be successfully applied to the establishment of coordinate systems for organs of varying degrees of morphological complexity and the subsequent quantitative analysis of 3D cellular parameters. We provided evidence for a preferential increase in cell length during axial neck growth of the Marchantia archegonium. Moreover, we identified distinct cellular patterns possibly associated with important morphological features of the intricately folded *U. gibba* trap. These data reveal that 3DCoordX has broad applicability and eliminates the need to work with multiple different pipelines when analyzing the cellular architecture of organs in 3D. The general strategies and computational methods put forward in this work will greatly reduce the time required for the spatial analysis of cellular parameters so central to various approaches, such as computational modeling of morphogenesis or comparative morphometry of specimens from different genotypes.

## Materials and Methods

### Plant work and lines

*Arabidopsis thaliana* (L.) Heynh. var. Columbia (Col-0) was used as a wild-type strain. Plants were grown on soil as described earlier [64]. The pKAN1::KAN1:2xGFP construct [38] and the pCUC3::CFP line [52] were gifts from Marcus Heisler and Nicolas Arnaud, respectively. Wild-type plants were transformed with the pKAN1::KAN1:2xGFP construct using Agrobacterium strain GV3101/pMP90 [65] and the floral dip method [66]. Transgenic T1 plants were selected on Glufosinate (Basta) (10 μg/ml) plates and transferred to soil for further inspection. *Marchantia polymorpha* of the BoGa ecotype was grown on half-strength Gamborg’s B5 medium under long day conditions (16L:8D) at 22°C. For induction of reproductive structures, plants were grown under 60 µmol white light supplemented with far red light (730 nm) on half-strength Gamborg’s B5 medium supplemented with 1 % glucose and 14 g/L agarose [67]. Gametangiophores appeared after 4–6 weeks.

### Clearing and staining of tissue samples

Treatment of ovules of the pKAN1::KAN1:2xGFP and the pCUC3::CFP lines was done as described in [68] and [27]. Tissue was fixed in 4% paraformaldehyde in PBS followed by two washes in PBS before transfer into the ClearSee solution (xylitol (10%, w/v), sodium deoxycholate (15%, w/v), urea (25%, w/v), in H2O) [69]. Clearing was done at least overnight or for up to two to three days. Cell wall staining with SR2200 (Renaissance Chemicals, Selby, UK) was performed as described in [70]. Cleared tissue was washed in PBS and then put into a PBS solution containing 0.1% SR2200 and a 1/1000 dilution of the nuclear stain TO-PRO-3 iodide (Thermo Fisher Scientific) for 20 minutes. Tissue was washed in PBS for one minute, transferred again to ClearSee for 20 minutes before mounting in Vectashield antifade agent (Vector Laboratories, Burlingame, CA, USA). Marchantia archegoniophores were fixed for 1 week in 4% paraformaldehyde in PBS followed by two washes in PBS before transfer to ClearSee. Clearing was done for 4-7 days. Cell wall staining and subsequent clearing, washing and mounting steps were the same as for Arabidopsis ovules. Archegonia were dissected in Vectashield mounting medium (Vector Laboratories, Burlingame, CA, USA).

### Microscopy and image acquisition

Confocal laser scanning microscopy of ovules stained with SR2200 and TO-PRO-3 iodide was performed on an upright Leica TCS SP8 X WLL2 HyVolution 2 (Leica Microsystems) equipped with GaAsP (HyD) detectors and a 63x glycerol objective (HC PL anterior-posteriorO CS2 63x/1.30 GLYC, CORR CS2). Scan speed was at 400 Hz, the pinhole was set to 0.6 Airy units, line average between 2 and 4, and the digital zoom between 1 and 2. For z-stacks, 12-bit images were captured at a slice interval of 0.24 μm with voxel size of 0.125 μm x 0.125 μm x 0.24 μm. Laser power or gain was adjusted for z compensation to obtain an optimal z-stack. Images were adjusted for color and contrast using Adobe Photoshop 2021 (Adobe, San Jose, USA) or MorphographX [14] software. Image acquisition parameters for the pKAN1::KAN1:2xGFP line were the following: SR2200; 405 diode laser 0.10%, HyD 420–480 nm, detector gain 10. 2xGFP; 488 nm Argon laser 2%, HyD 525-550 nm, detector gain 100. TO-PRO-3; 642 nm White Laser 2%, HyD 660–720 nm, detector gain 100. In each case sequential scanning was performed to avoid crosstalk between the spectra. Image acquisition parameters for the pCUC3::CFP line were the following: SR2200; 405 diode laser 0.10%, HyD 420–480 nm, detector gain 10. CFP; 514 nm Argon laser 2%, HyD 525-550 nm, detector gain 100. TO-PRO-3; 642 nm White Laser 2%, HyD 660–720 nm, detector gain 100. In each case sequential scanning was performed to avoid crosstalk between the spectra. Imaging conditions for the Marchantia archegonia stained with SR2200 and TO-PRO-3 iodide were the same as for the ovules. The late stage Marchantia archegonium was imaged using the same 63x glycerol objective and a tilescan of 8 tiles. For z-stacks of the older specimen, 8-bit images were captured at a slice interval of 0.33 μm with voxel size of 0.126 μm x 0.126 μm x 0.33 μm; for z-stack of the younger 12-bit images were captured at a slice interval of 0.33 μm with voxel size of 0.127 μm x 0.127 μm x 0.33 μm. Tiles were stitched and merged to form the final 3D image stack of the organ in Leica Application Suite X data processing software (LASX v3.5.7.23225). The early stage archegonium was imaged without tile scan.

### Datasets and 3D cell segmentation

The dataset encompassing the segmented wild-type 3D digital ovules was described earlier [27]. The two z-stacks of Marchantia archegonia were 3D cell segmented using the PlantSeg pipeline [17]. The z-stack of the *Utricularia gibba* trap was obtained from a fixed and modified pseudo-Schiff-stained [71] specimen [59]. 3D cell segmentation was performed using the PlantSeg-MorphoGraphX hybrid method as described in [27]. In all instances generation of cell surface meshes and cell type labeling was performed with MorphoGraphX.

### Software

The MorphographX software was used for the generation of cell surface meshes, cell type labeling, and the analysis of 3D cellular features [14]. It can be downloaded from its website (https://www.mpiz.mpg.de/MorphoGraphX). The 3DCoordX toolbox is integrated as an add-on in MorphoGraphX 2.0. A detailed user manual is provided in the supplement. The PlantSeg pipeline [17] was used for 3D cell boundary prediction and segmentation. The software can be obtained from its Github repository (https://github.com/hci-unihd/plant-seg).

## Acknowledgements

We thank Marcus Heisler and Nicolas Arnaud for the pKAN1::KAN1:2xGFP and pCUC3::CFP lines, respectively. We also thank Peter Schroeder and Claus Schwechheimer for providing *Marchantia polymorpha* female gametophyte samples. We thank members of the Schneitz lab for helpful discussions and Adam Runions for discussions on growth alignment graphs. We further thank Enrico Coen for insightful comments. We acknowledge support by the Center for Advanced Light Microscopy (CALM) of the TUM School of Life Sciences. This work was funded by the German Research Council (DFG) through grants FOR2581 (TP7) to KS, (TP8) to RS, (TP9) to MT and a Max Planck Society core grant to MT. Work by KL was funded by the Biotechnology and Biological Sciences Council (BBSRC) through grants BBS/E/J/00000152, BBS/E/J/000PR9787, and BBS/E/J/000CA517.

## Authors’ contributions

AV, SS, RS and KS designed the study. AV, SS, RT, TAM, and KL performed the experiments. AV, SS, RT, TAM, KL, MT, RS and KS interpreted the results. MT, RS and KS secured funding. KS wrote the paper with comments from all authors. All authors read and approved the final manuscript.

---

**Fig S1.**
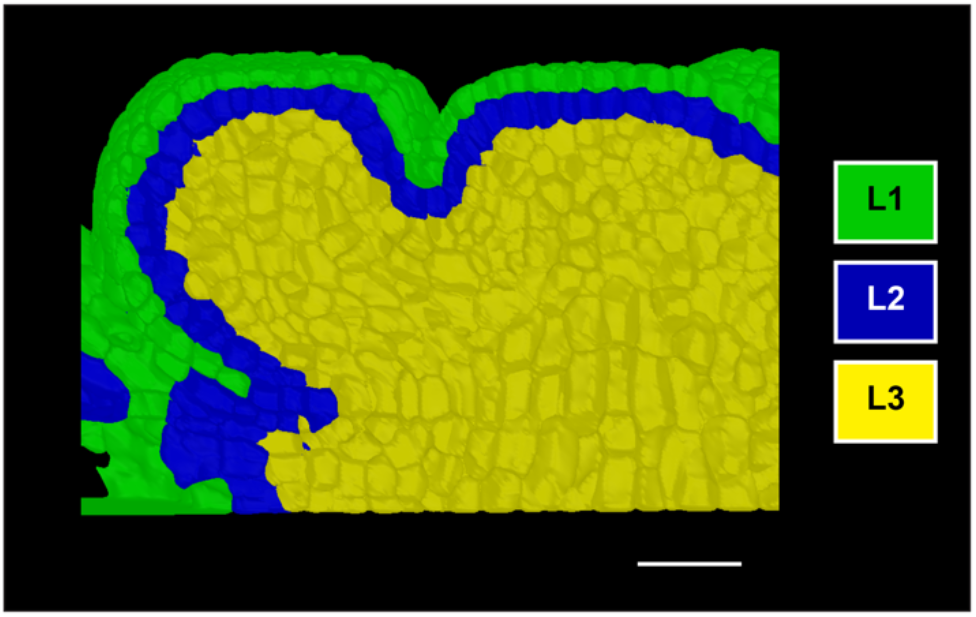
Automatic cell layer detection in the Arabidopsis shoot apical meristem. The outside wall area ratio approach was applied to automatically identify the three tissue layers. Layers are labelled with respective colors. Scale bar 20 µm.

**Fig. S2.**
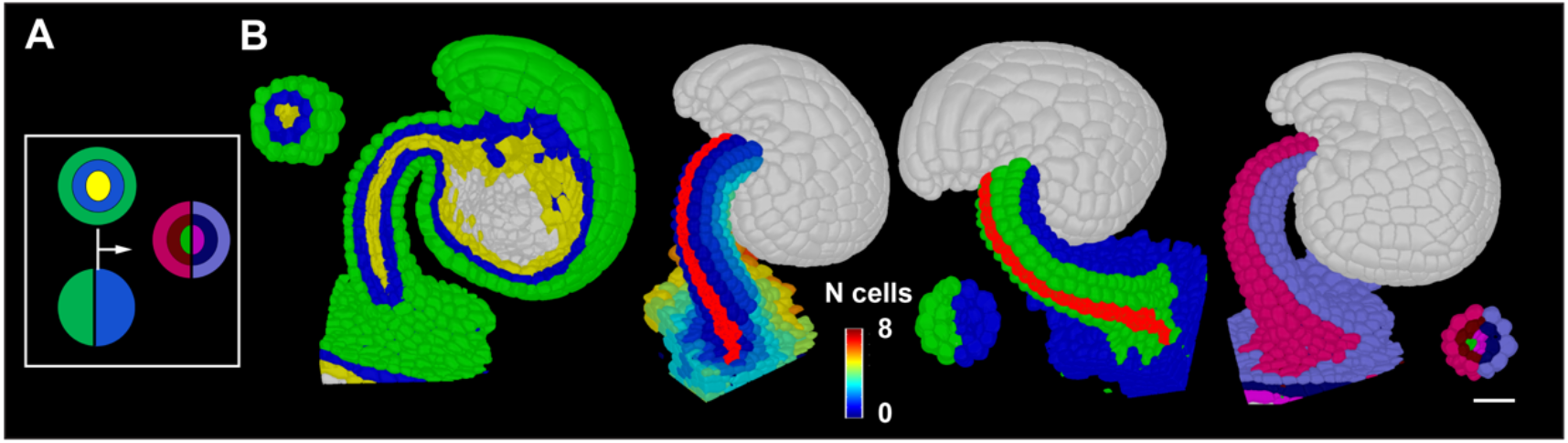
Separation of funiculus into radial and anterior and posterior domains. A cartoon demonstrating the principle for the separation into radial L1, L2, L3 layers and anterior-posterior domains. (A) Results of L1, L2 and L3 tissue labels are combined with anterior and posterior labels to form the final tissue labels that separate individual radial layers and anterior-posterior tissues. (B) Figure demonstrating the step-by-step procedure for the detection of the anterior and posterior regions of the Arabidopsis funiculus. From left to right; first panel: L1, L2 and L3 labelled Arabidopsis ovule, a transverse section is shown at the top left; second panel: cells at roughly anterior midline are selected as origin (highlighted in red) and cell distance from these cells to other cells in funiculus are computed. Essentially the cell distance here represents the number of cells a cell is away from the anterior midline cells; third panel: the heatmap of cell distances from the second image is binned into two resulting in anterior and posterior tissue annotations; fourth panel: the results from first and third panel are merged to form the final tissue label that combines the information for L1, L2, L3 and anterior and posterior. This allows for example the distinction of anterior L1 and posterior L1 tissue. Scale bar 20 µm.

